# Transgenic augmentation of erythroferrone in mice ameliorates anemia in adenine-induced chronic kidney disease

**DOI:** 10.1101/2024.12.06.627111

**Authors:** Brian Czaya, Joseph D. Olivera, Moya Zhang, Amber Lundin, Christian D. Castro, Grace Jung, Elizabeta Nemeth, Tomas Ganz

## Abstract

Anemia is a common and disabling complication of chronic kidney disease (CKD). Current therapies can be burdensome, and full correction of anemia is limited by cardiovascular side effects. New approaches that may offer additional therapeutic options are needed. We explored the anti-anemic effects of erythroferrone, an erythroid hormone that induces iron mobilization by suppressing the master iron-regulatory hormone hepcidin. In a preclinical murine model of adenine-induced CKD, transgenic augmentation of erythroferrone mobilized iron, increased hemoglobin concentrations by approximately 2 g/dl, and modestly improved renal function without affecting systemic or renal inflammation, fibrosis, or markers of mineral metabolism. This study supports the concept that therapeutic augmentation of erythroferrone is a promising approach for alleviating CKD-associated anemia.

## Introduction

Chronic kidney disease (CKD) affects more than 10% of the population in the United States (1). Anemia, often severe if untreated, is a major morbidity in CKD and affects most of the patients with advanced CKD (2). The pathogenesis of this anemia is multifactorial, with mechanisms including impaired production of erythropoietin, dysregulated iron homeostasis, and the suppressive effects of inflammation and uremic toxins on erythropoiesis (3). Although the combination of erythropoietin derivatives and intravenous iron is effective in treating anemia (4), the logistical burdens for those patients who are not receiving regular hemodialysis have interfered with wider adoption of these therapies (5). Moreover, treatment with erythropoietin derivatives increases cardiovascular complications, and therefore is subject to limits that do not allow complete reversal of anemia. Orally administered prolyl hydroxylase inhibitors are also effective for treating anemia and more convenient for outpatient treatment but they have not demonstrated improved cardiovascular risks compared to erythropoietin and IV iron (4). Additional treatment options are clearly needed (6).

Erythroferrone (ERFE) is a hormone produced by red cell precursors in the marrow and secreted into the blood stream (7). Its main documented effect is the transcriptional suppression of the hepatic iron-regulatory hormone hepcidin. Decreased hepcidin concentrations in blood then allow increased intestinal iron absorption and increased mobilization of iron from stores. In effect, physiologic augmentation of erythroferrone, such as occurs after blood loss or the administration of erythropoietin, facilitates the provision of iron for the increased requirements of intensified erythropoiesis.

To understand the effects of chronically augmented ERFE production, we generated transgenic mice that selectively overexpressed ERFE in red blood cell precursors, yielding three transgenic strains with graded overexpression of ERFE (8). We observed a dose-dependent effect of ERFE on iron loading, with the highest overexpressors, line-H, developing considerable systemic iron overload and a blood hemoglobin increase of 1 g/dl.

In the current study, we explored whether ERFE augmentation could be used as a treatment for CKD-associated anemia. We used the mouse model of adenine-induced nephropathy, in which excessive amounts of dietary adenine are administered, and converted *in vivo* by xanthine oxidase to 2,8-dihydroxyadenine which precipitates in the urine causing tubulointerstitial kidney disease (9). A human genetic disorder, adenine phosphoribosyl transferase deficiency, causes a similar disease in humans (10). This mouse model recapitulates many characteristics of human CKD including anemia, inflammation and iron restriction (9, 11, 12).

## Results

In an initial pilot experiment, we exposed line-H (high-expressing) ERFE-transgenic mice (8) and their wild-type littermates to 0.2% adenine diet containing 100 ppm iron for 8 weeks (Fig. 1A). As expected, both WT and TG mice developed anemia, appeared ill and lost weight. However, after 8 weeks of adenine, TG compared to WT mice had higher hemoglobin (Hb) concentrations by 2 g/dl, as well as higher RBC count and mean corpuscular hemoglobin (MCH) (Fig. 1 B-E). Although these findings are informative, we cannot differentiate the contribution of preexisting iron excess and higher hemoglobin in TG mice at the start of adenine diet versus improved iron availability during the course of disease in this model, as TG mice have liver iron concentrations 4.2-times higher than WT, and have 1 g/dl higher Hb concentrations by the age of 6 weeks (8).

**Figure 1.**
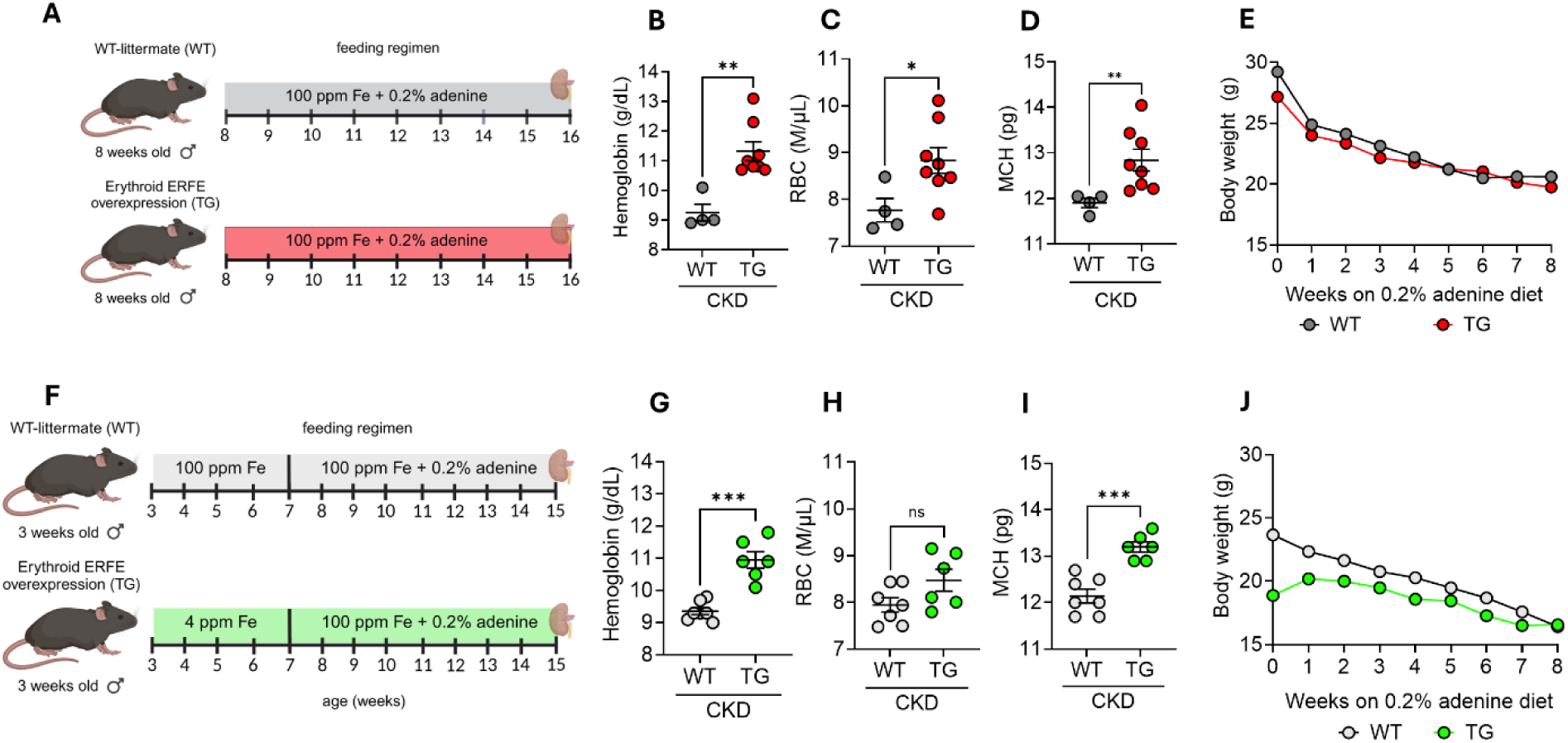
ERFE augmentation ameliorates CKD-associated anemia. (**A**) Pilot experimental design: 8-week-old WT-littermate (WT; n = 4) and ERFE-overexpressing transgenic (TG; n = 7) mice were fed 100 ppm iron (Fe) diet with 0.2% adenine for 8 weeks then analyzed. (**F**) Final experimental design: Mice were weaned at 3 weeks, then WT (n = 7) mice placed on 100 ppm Fe diet and TG (n = 6) placed on 4 ppm Fe diet to prevent iron accumulation prior to the initiation of adenine diet. At 7 weeks of age WT and TG mice were placed on 0.2% adenine diet with 100 ppm Fe for 8 weeks then analyzed. For both experiments: (**B, G**) blood hemoglobin concentration, (**C, H**) RBC count, (**D, I**) mean corpuscular hemoglobin (MCH). (**E, J**) Mice were weighed weekly. Data are mean ± SEM, analyzed by unpaired-*t*-test with Welch’s correction (two-tailed). ***P <.001; **P <.01; *P <.05; ns = non-significant.

To develop a more clinically-relevant model, we noted that at weaning at 3 weeks of age TG mice had not yet developed iron overload, so that TG compared to WT mice had similar body weights, erythrocyte parameters and slightly lower liver iron concentrations (LIC) (Fig. S1A-F). We observed that placing 3-week-old TG mice on an essentially iron-free (4 ppm) diet for 4 weeks (until the intended start of adenine diet) prevented iron accumulation, and these 7-week-old TG mice had even lower Hb concentrations, RBC count, MCH, liver iron and serum iron than their littermate WT mice (Fig. S1G-L).

To induce CKD, parallel cohorts of 7-week-old iron-depleted TG mice and their 7-week-old WT littermates were placed on 0.2% adenine diet containing 100 ppm Fe (Fig. 1F). Despite TG iron depletion regimen causing greater anemia and iron deficiency at the onset of adenine diet, after 8 weeks on adenine diet, TG mice had higher Hb (by approx. 2 g/dl) and higher MCH than WT mice (Fig. 1 G-I). Remarkably, the change in Hb between the pre-and post-adenine groups was-4.7 g/dL for WT mice and +2.8 g/dL for TG mice (compare Fig. S1H and Fig. 1G). Furthermore, despite starting at a lower body weight, TG mice ended up weighing the same as WT mice (Fig. 1J, Fig. 2A and Supplemental Table I).

**Figure 2.**
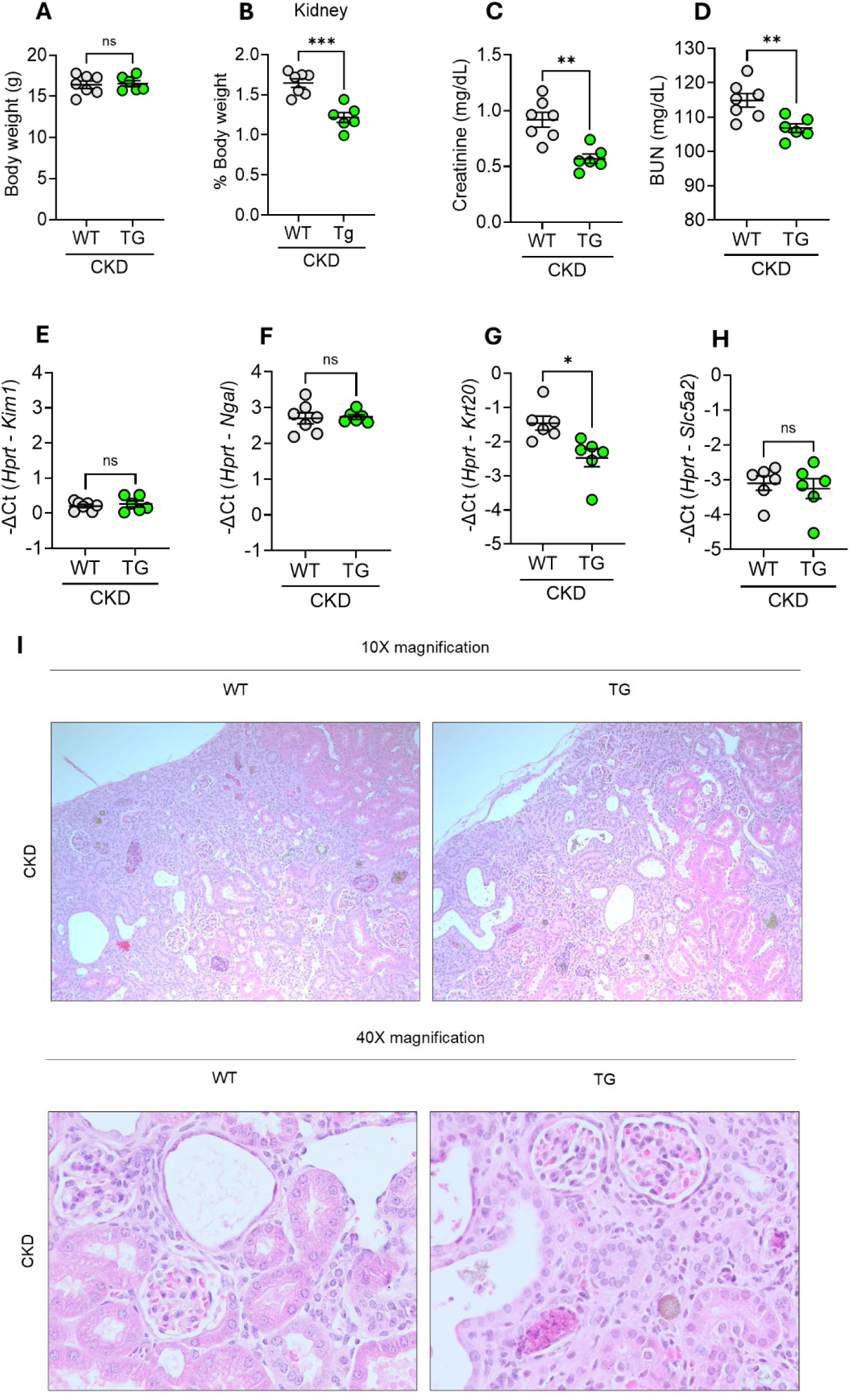
ERFE augmentation improves kidney function but does not alter injury markers in adenine-induced CKD. WT and TG mice from Figure 1F (n=6-7 mice/group) were analyzed after 8 weeks on adenine diet. (**A**) Body weight, (**B**) kidney weight, (**C**) serum creatinine, (**D**) blood urea nitrogen (BUN). Quantitative PCR (qPCR) analysis of (**E**) *Kim1*, (**F**) *Ngal,* (**G**) *Krt20* and (**H**) *Slc5a2* expression in kidney tissue. (**I, J**) Representative H&E-stained kidney sections from WT and TG mice after 8 weeks on adenine diet (10x and 40x magnification, respectively). Data are mean ± SEM, analyzed by unpaired-*t*-test with Welch’s correction (two-tailed). ***P <.001; **P <.01; *P <.05; ns = non-significant.

As we previously reported (8), the exposure to ERFE during development results in TG mice having slightly smaller kidneys and mild impairment of renal function at baseline. Indeed, after 8 weeks of adenine diet, TG mice still exhibited smaller kidneys than WT mice (Fig. 2B). However, while renal function was impaired in both WT and TG mice (higher BUN compared to normal ∼20-30 mg/dL (8)), both BUN and creatinine were lower in TG than WT mice, indicating significant amelioration of CKD by transgenic ERFE. The expression levels of certain renal injury markers (*Kim1*, *Ngal* and SGLT2 (*Slc5a2*)) were similar in WT and TG mice, whereas *Krt20* was lower in TG, consistent with some degree of protection from kidney injury (Fig. 2F-I). The histological appearance of kidney sections was similar in WT and TG mice, and consistent with adenine-induced tubulointerstitial damage.

To investigate the mechanism of anemia amelioration by transgenic ERFE, we examined systemic iron parameters (Fig. 3). After 8 weeks of adenine diet, TG mice had higher liver iron concentrations (reflective of iron stores), higher serum iron and non-significantly lower splenic iron than WT mice (Fig 3A-C). This indicates that TG mice had increased iron absorption (reflected by LIC) and possibly increased iron mobilization from red pulp macrophages (reflected by spleen iron). Kidney iron did not significantly differ between TG and WT mice. Serum hepcidin and hepatic hepcidin expression were lower in TG than in WT mice, despite higher LIC, and the ratio of hepatic hepcidin expression to LIC was markedly lower in TG vs WT mice, consistent with the documented hepcidin-suppressive effect of ERFE (7, 8). Lower hepcidin concentrations and improved iron availability for erythropoiesis in TG compared to WT mice would be expected to increase Hb concentrations and RBC hemoglobinization, as was observed (Fig. 1).

**Figure 3.**
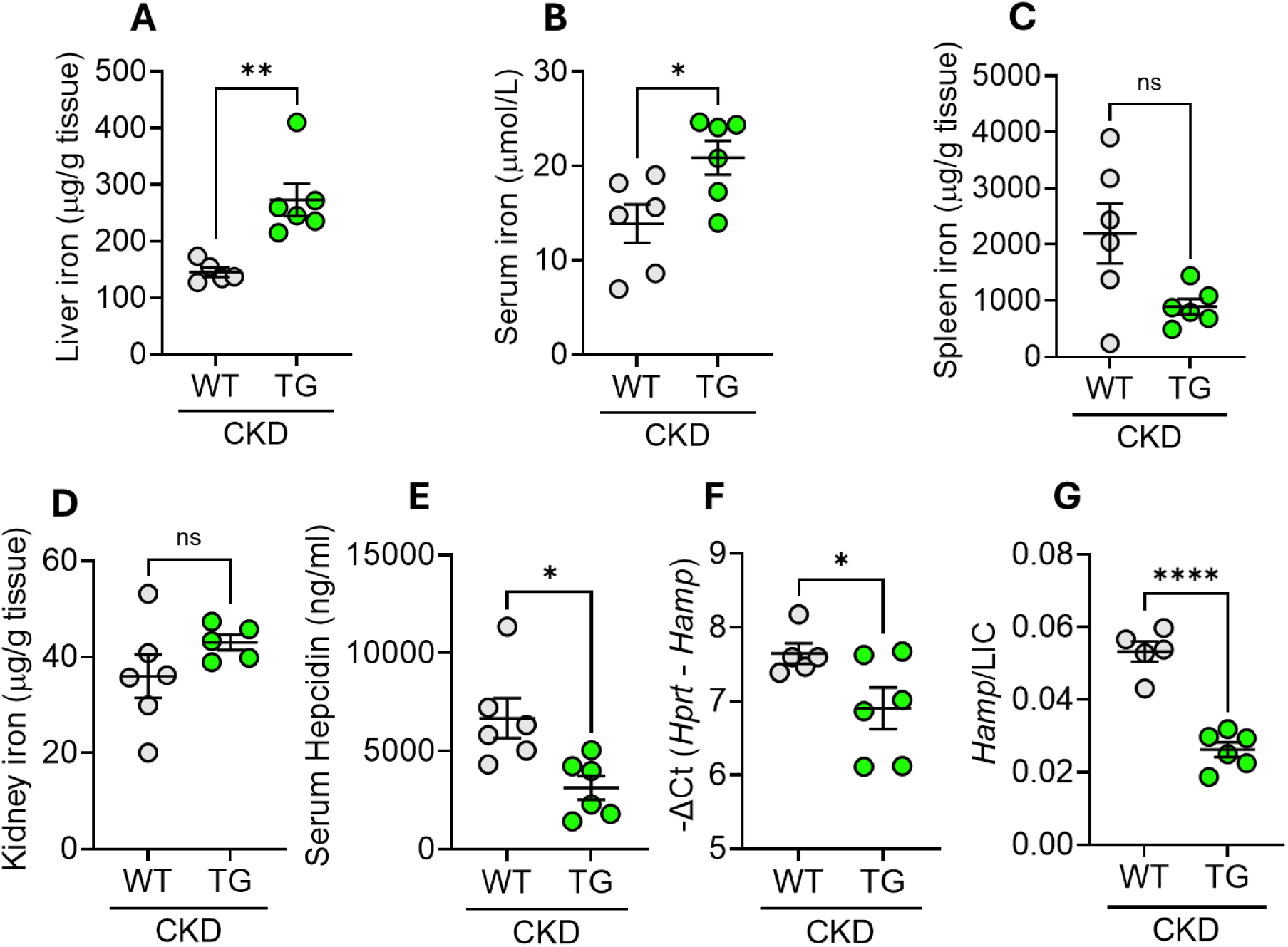
ERFE augmentation lowers hepcidin levels to enhance iron mobilization in adenine-induced CKD. WT and TG mice from Figure 1F (n=6-7 mice/group) were analyzed. (**A**) LIC, (**B**) serum iron, (**C-D**): tissue iron concentrations in (**C**) spleen and (**D**) kidney, (**E**) serum hepcidin levels, (**F**) liver hepcidin mRNA expression, and **(G)** liver hepcidin mRNA/LIC. Data are mean ± SEM, analyzed by unpaired-*t*-test with Welch’s correction (two-tailed). ****P<.0001, **P <.01; *P <.05; ns = non-significant.

The effects of ERFE on kidney injury could be complex. On one hand, amelioration of anemia would be expected to improve oxygen delivery and benefit the energy-intensive functional and repair processes in the kidney. On the other hand, ERFE’s mechanism of action depends on inhibition of BMP signaling involved not only in the regulation of hepcidin transcription (13–15) but also in kidney development and repair (16–18).

We first examined the effects of transgenic ERFE on hypoxia-sensitive processes in adenine-CKD mice (Fig. 4). Serum VEGF was lower in TG than in WT mice, indicative of improved tissue oxygenation in TG mice. In the kidney at this advanced disease stage, the expression of *Vegfa, Gapdh, Angptl1* and *Epo* all trended lower in TG than in WT mice but the differences were small and not statistically significant.

**Figure 4.**
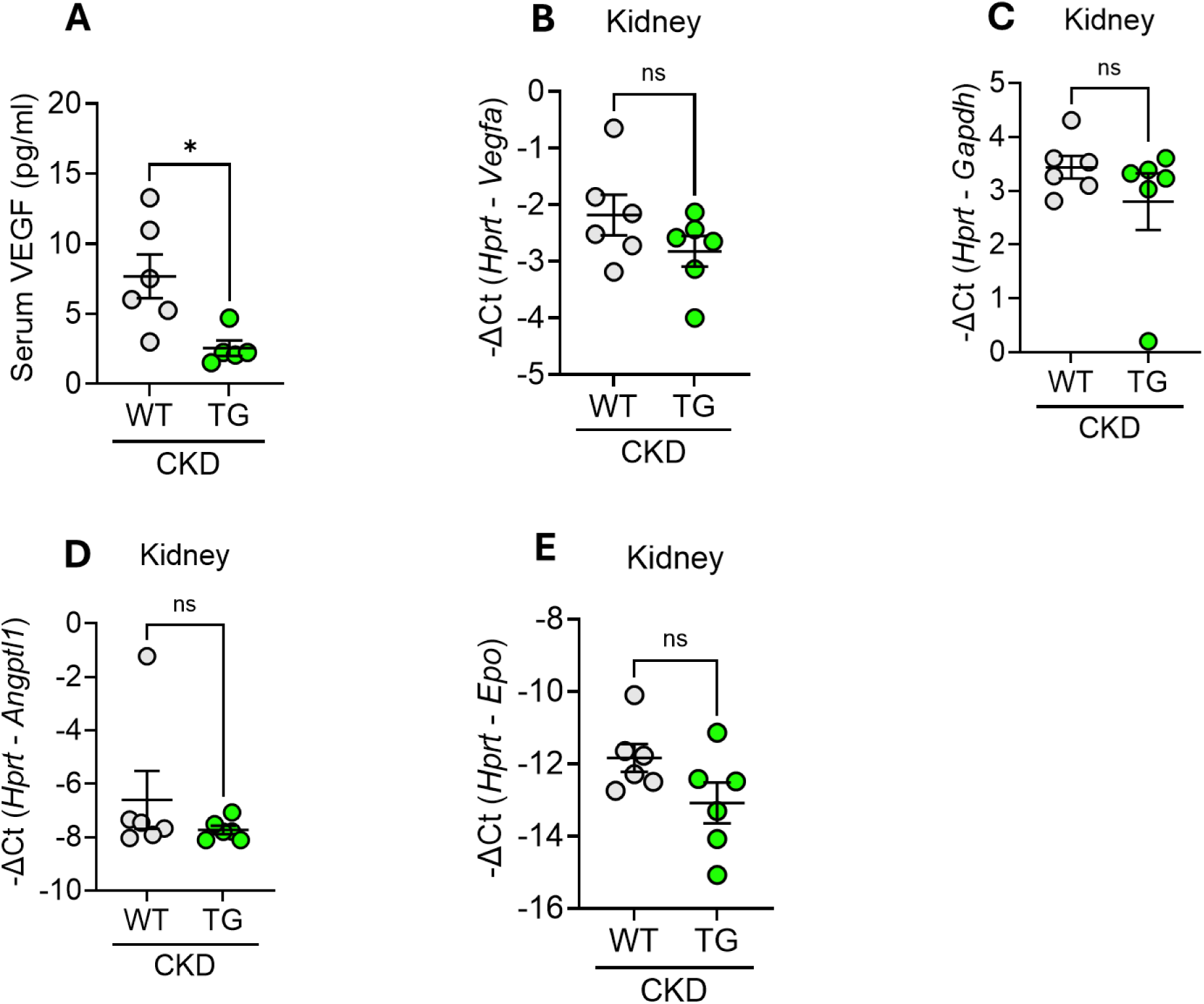
ERFE augmentation enhances systemic oxygenation without significantly impacting kidney oxygenation in adenine-induced CKD. WT and TG mice from Figure 1F (n=6-7 mice/group) were analyzed: (**A**) serum VEGF levels, **(B-E):** kidney mRNA concentrations by qRT-PCR (**B**) V*egfa*, (**C**) *Gapdh*, (**D**) *Angptl1* and (**E**) *Epo*. Data are mean ± SEM, analyzed by unpaired-*t*-test with Welch’s correction (two-tailed). *P <.05; ns = non-significant.

We next analyzed the expression of several genes affected by BMP signaling in the kidney (Fig. 5). Among these, the expression of *Id1, Id2, Id4* as well as *Bmp2* and *Bmp7* was similar in TG and WT mice but *Smad7* was lower in TG mice. The expression of renal fibrosis markers could potentially be affected by BMP signaling, but levels of *Acta2* (alpha smooth muscle actin), *Col1a1* and *Col3a1* (type I and III collagen a1 chains), *Tgfb* (TGF-β1) and *Fn1* (fibronectin) were not significantly changed. Mason’s trichrome staining of kidney tissue sections also had a similar appearance (Fig. 6). In the aggregate, although there was a detectable suppression of a sensitive BMP signaling-related marker, there was no appreciable effect on renal fibrosis.

**Figure 5.**
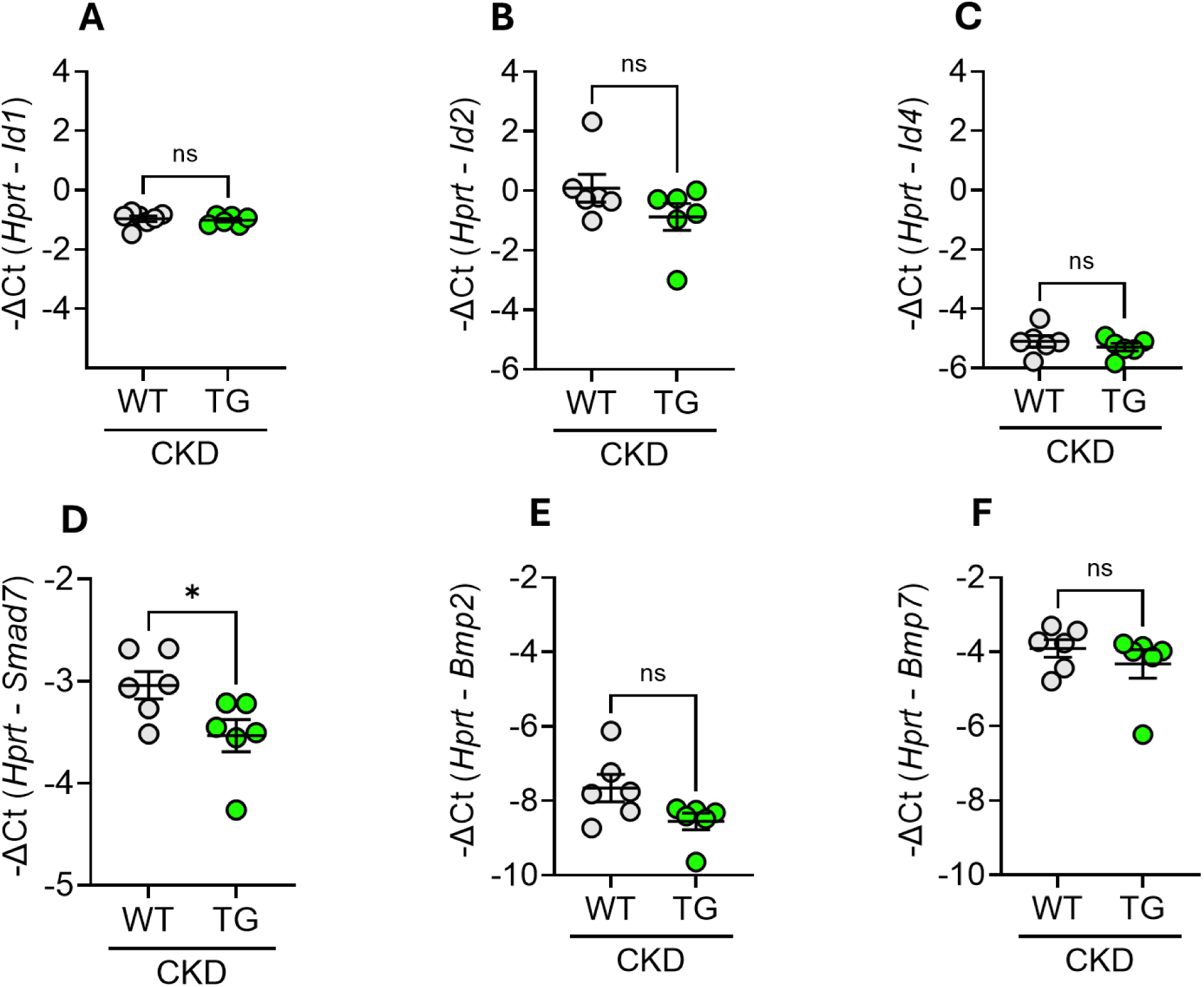
ERFE augmentation results in unchanged or mildly suppressed kidney BMP signaling in adenine-induced CKD. WT and TG mice from Figure 1F (n=6-7 mice/group) were analyzed by qRT-PCR of kidney tissue for (**A**) *Id*1, (**B**) *Id2*, (**C**) *Id4*, (**D**) *Smad7*, (**E**) *Bmp2* and (**F**) *Bmp7*. Data are mean ± SEM, analyzed by unpaired-*t*-test with Welch’s correction (two-tailed). *P <.05; ns = non-significant.

**Figure 6.**
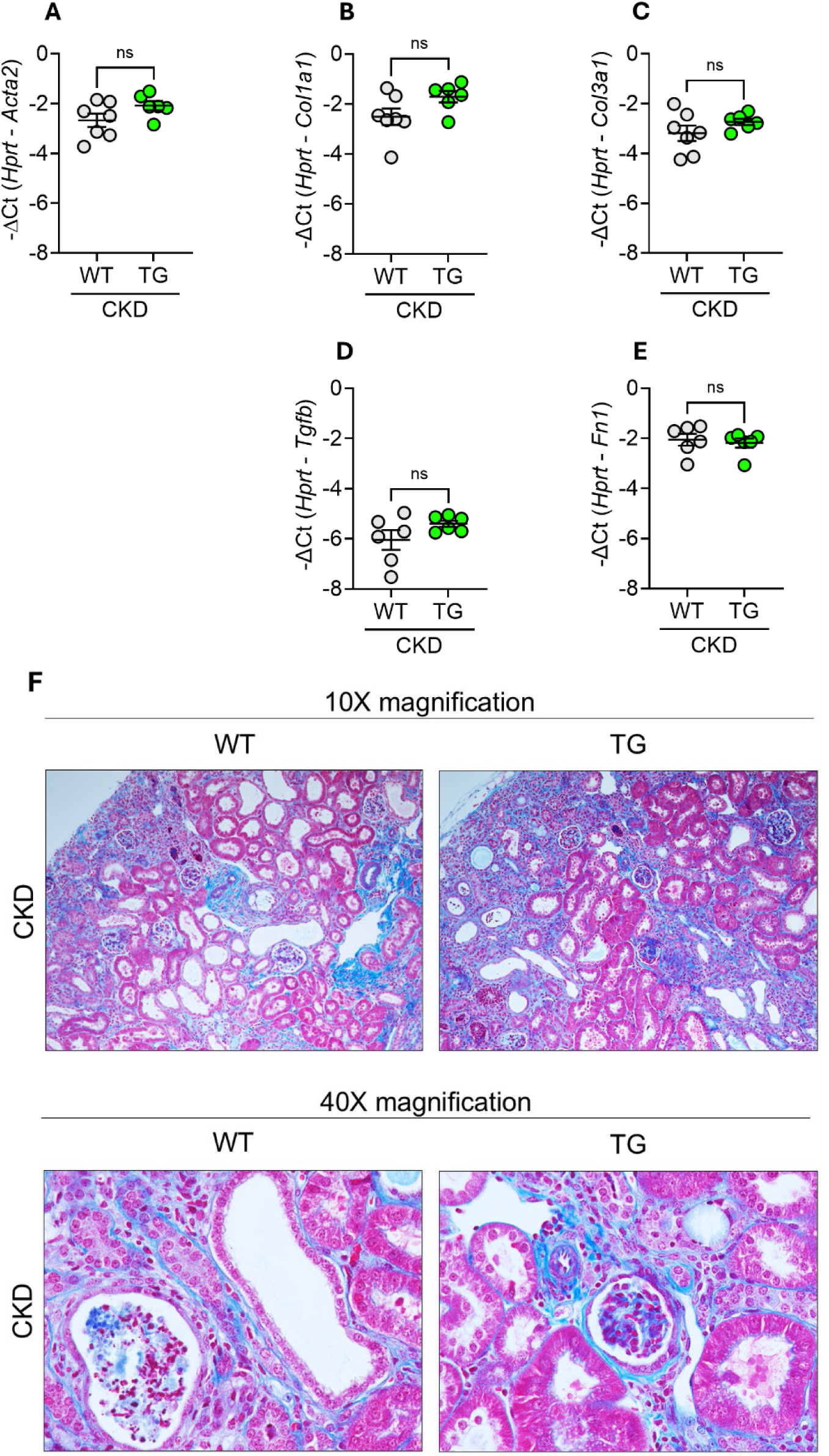
ERFE augmentation does not affect kidney fibrosis in adenine-induced CKD. In WT and TG mice from Figure 1F (n=6-7 mice/group), kidney tissues were analyzed by qRT-PCR for (**A**) *Acta2*, (**B**) *Col1a1*, (**C**) *Col3a1*, (**D**) *Tgfb1* and (**E**) *Fn1*. (**F**) Representative Masson trichrome-stained kidney sections from WT and TG mice after 8 weeks on adenine diet (10x and 40x magnification). Data are mean ± SEM, analyzed by unpaired-*t*-test with Welch’s correction (two-tailed). ns = non-significant.

Inflammation is an important driver of both anemia (19) and kidney injury (20). We therefore examined whether transgenic ERFE affected systemic and organ inflammation elicited by 2,8- dihydroxyadenine crystals in the kidney and by the subsequent impairment of renal function. We found no significant differences between TG and WT mice in plasma concentrations of key inflammatory cytokines TNFα, IL1β, IL6, IL1α, IFNγ or IL10 (Fig. 7). The expression of the acute phase reactant and hepatic inflammatory marker SAA1 and that of renal cytokines IL6 and IL1β were similar between WT and TG mice with adenine-induced CKD, while renal expression of TNFa was mildly increased in TG compared to WT mice (Fig. 8). Overall, ERFE overexpression did not have a substantial effect on systemic or renal inflammation.

**Figure 7.**
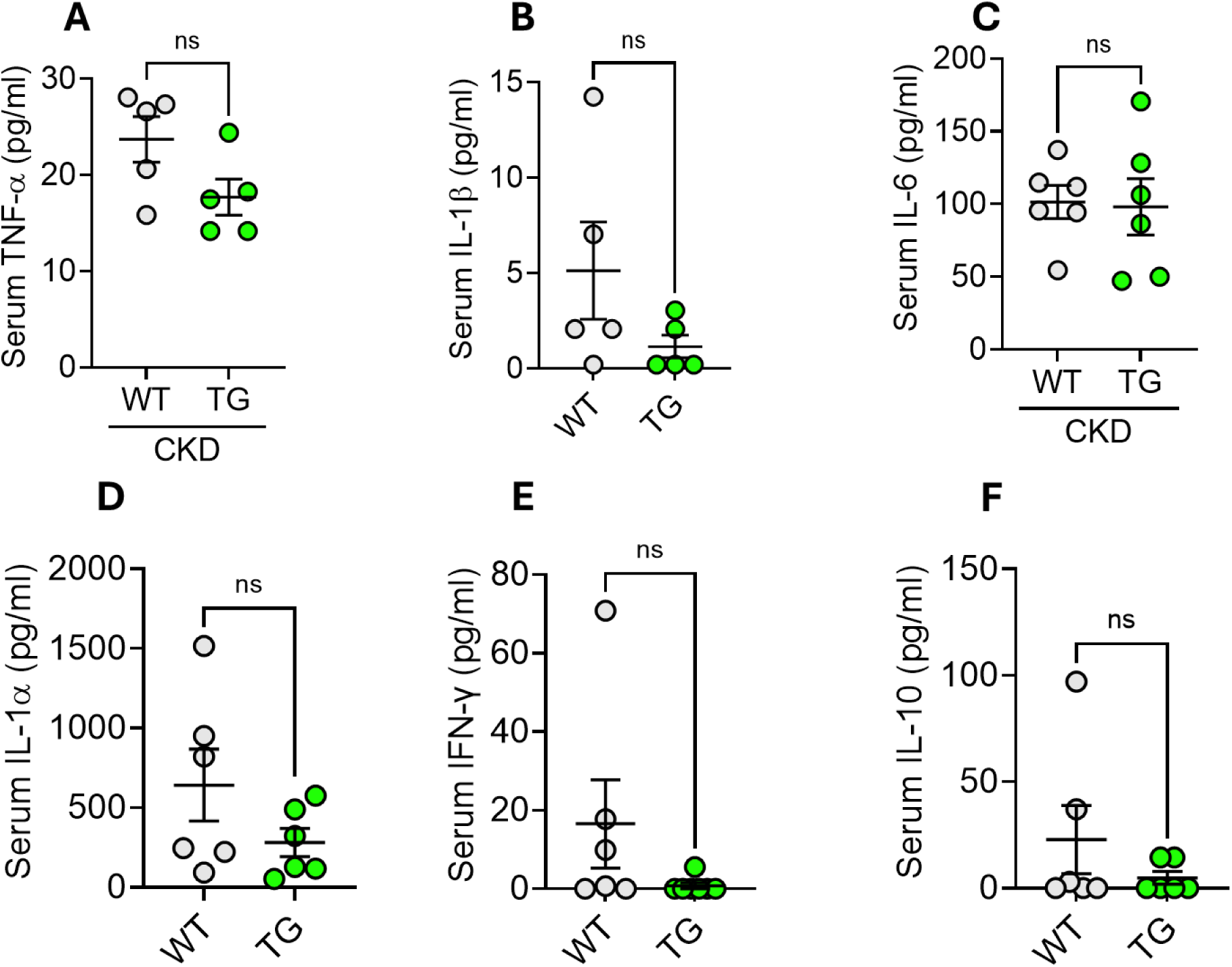
ERFE augmentation does not impact systemic inflammation in adenine-induced CKD. WT and TG mice from Figure 1F (n=6-7 mice/group) were analyzed by serum 32-plex cytokine/chemokine assay for (**A**) TNFα, (**B**) IL-1β, (**C**) IL-6, (**D**) IL-1α, (**E**) IFNγ, and (**F**) IL-10. Data are mean ± SEM, analyzed by unpaired-*t*-test with Welch’s correction (two-tailed). ns = non-significant.

**Figure 8.**
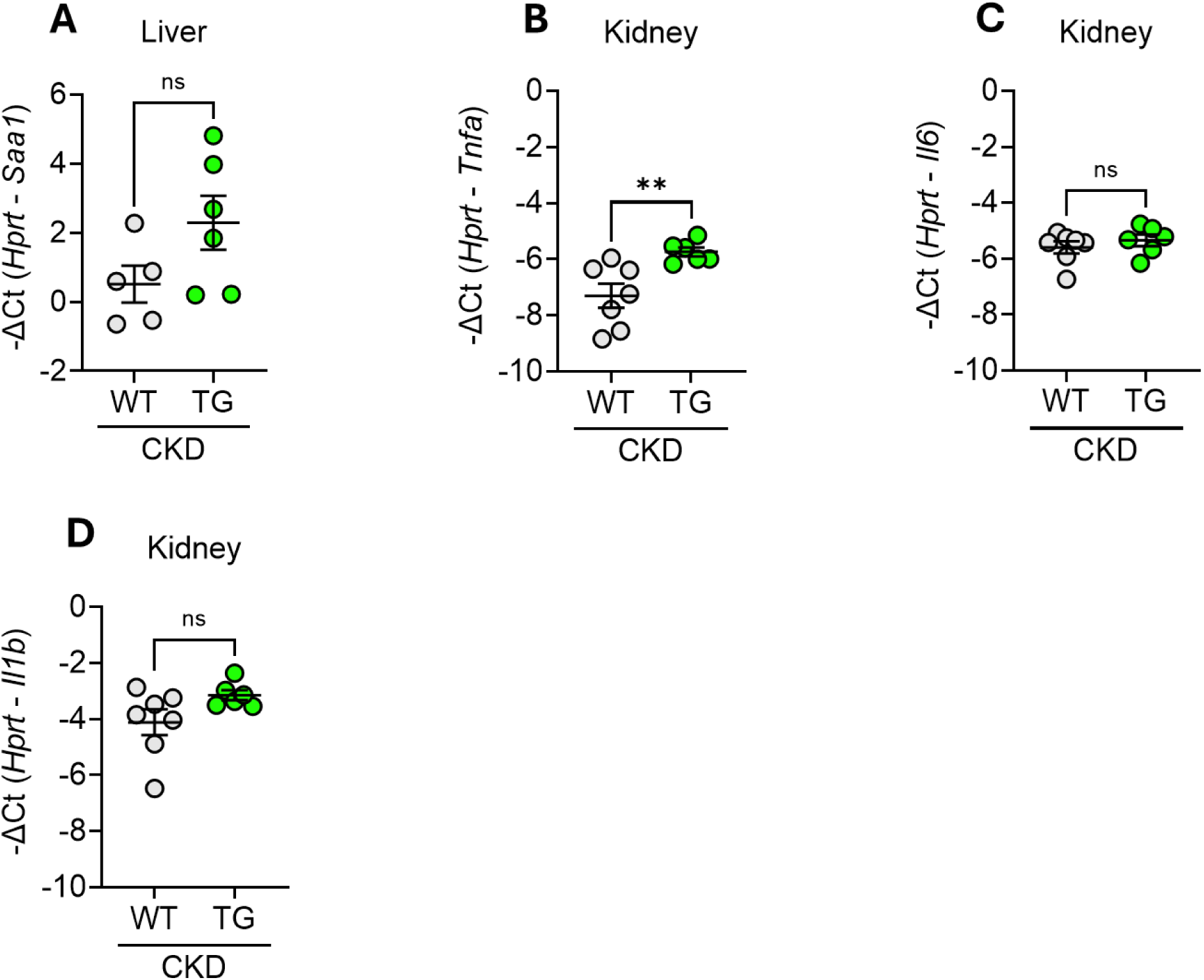
ERFE augmentation does not substantially change mRNA markers of liver or kidney inflammation in adenine-induced CKD. WT and TG mice from Figure 1F (n=6-7 mice/group) were analyzed by qRT-PCR for (**A**) *Saa1* expression in liver tissue, and (**B-D**) *Tnfa, Il6*, and *Il1b* expression in kidney tissue. Data are mean ± SEM, analyzed by unpaired-*t*-test with Welch’s correction (two-tailed). **P <.01; ns = non-significant.

Dysregulated mineral metabolism in CKD is closely linked to iron dyshomeostasis and contributes to adverse patient outcomes, including bone diseases and cardiovascular disease, in part through greatly increased FGF23 levels (21, 22). Linked to FGF23 excess, left ventricular hypertrophy (LVH) is a common and serious complication of CKD (23, 24). Accordingly, we analyzed (Fig. 9) the effect of ERFE overexpression on serum phosphate and iFGF23 (biologically active form) concentrations, as well as markers of mineral metabolism including *Slc34a1* and *Slc34a3* (encoding sodium-dependent phosphate transporters 2A and 2C also known as NaPi-2a and-2c), *Cyp27b1* and *Cyp24a1* (encoding the enzymes that respectively generate and break down the active form of vitamin D, 1,25-dihydroxyvitamin D3) and *Kl* (encoding Klotho, the renal coreceptor for iFGF23). No significant effects of transgenic ERFE overexpression were observed, except for a small increase in the expression of the phosphate transporter *Slc34a3* that mediates phosphate reabsorption in the renal tubular brush border. This increase may reflect the improved kidney function in TG mice (Figure 2).

**Figure 9.**
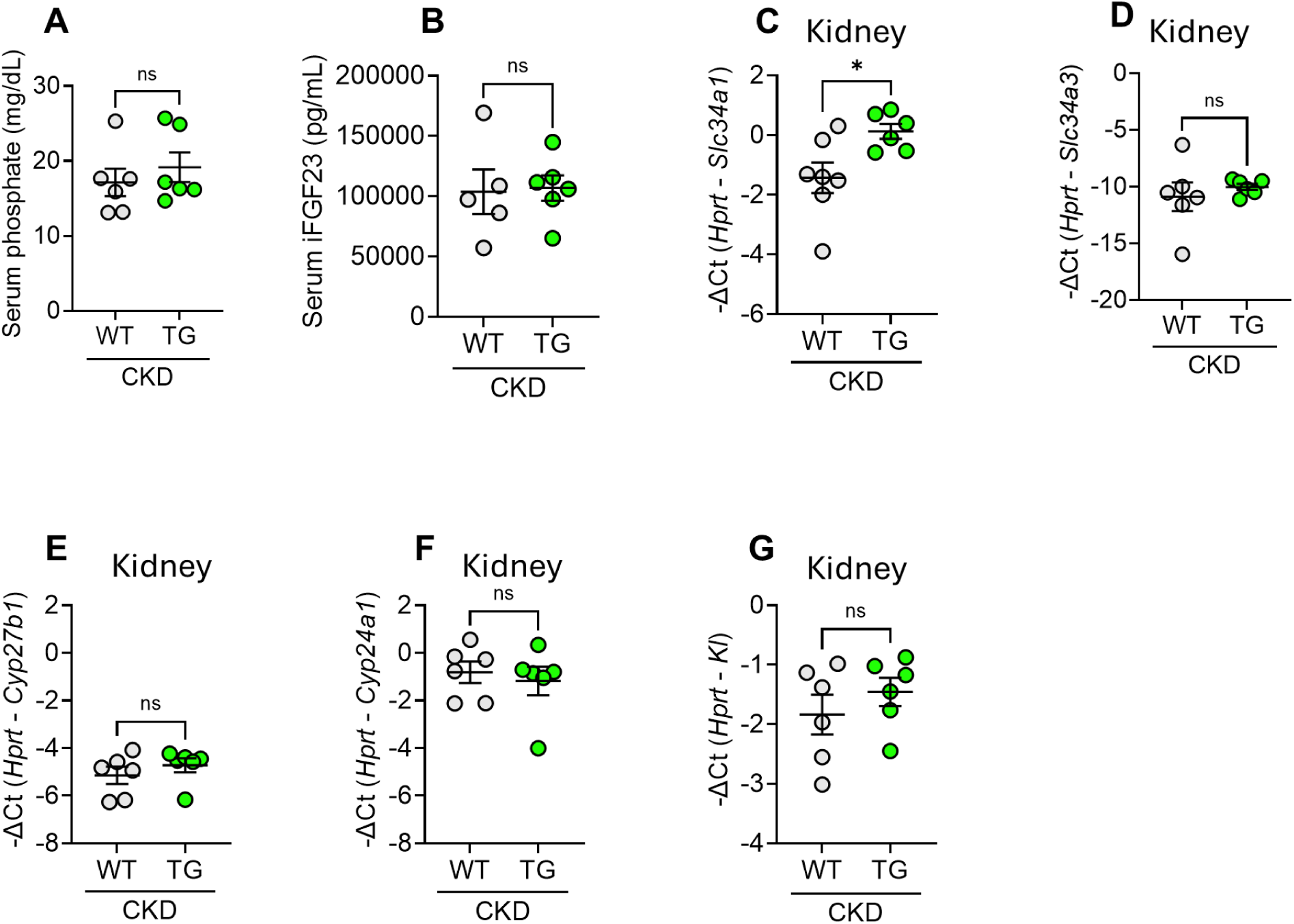
ERFE augmentation does not alter markers of mineral metabolism in adenine-induced CKD. WT and TG mice from Figure 1F (n=6-7 mice/group) were analyzed. Measured parameters include (**A**) serum phosphate, (**B**) serum intact iFGF23, and qPCR analysis of kidney tissue for (**C-G**) *Slc34a1* and *Slc34a3 (*encoding sodium-dependent phosphate transporters 2A and 2C), *Cyp27b1* and *Cyp24a1 (*encoding the enzymes that respectively generate and break down the active form of vitamin D, 1,25-dihydroxyvitamin D3) and *Kl* (encoding Klotho, the renal coreceptor for iFGF23). Data are mean ± SEM, analyzed by unpaired-*t*-test with Welch’s correction (two-tailed). *P <.05; ns = non-significant.

## Discussion

In this proof-of-concept study in mice, we showed that ERFE overexpression improved anemia in the dietary-adenine-induced model of CKD. At the end of the study, Hb was 2 g/dl higher in TG compared to WT mice, and the difference was even greater when Hb change is considered relative to the baseline pre-adenine group (-4.7 g/dL for WT mice and +2.8 g/dL for TG mice). These Hb improvements represent clinically significant increases. We noted no adverse effects in TG compared to WT mice during CKD progression. At the endpoint of the study, kidney function as measured by serum creatinine and BUN was somewhat better in TG than in WT mice, as reflected by multiple markers of kidney function and kidney injury (Figs. 2 and 9).

As shown in this study and a preceding report (8), the primary pathway impacted by transgenic overexpression of ERFE is iron homeostasis. The effect of ERFE augmentation is to increase liver iron concentration (LIC) and reduce iron content in the spleen. These effects are attributable to ERFE’s well-documented ability to suppress hepcidin, by inhibiting hepatic BMP signaling (13–15), the key pathway regulating hepcidin transcription in hepatocytes (25). Hepcidin suppression results in enhanced dietary iron absorption and mobilization of iron from macrophage stores. Clinically, these findings suggest that in CKD patients ERFE could promote the availability of sequestered dietary or medicinal iron for erythropoiesis, and enhance erythropoietic response to therapeutic erythropoietin.

Apart from animal studies, genetic evidence in humans indicates that ERFE would also mobilize iron and stimulate erythropoiesis in humans, as human *ERFE* gene intronic variant rs13007705-T leads to increased transferrin saturation, Hb and MCH in genome-wide association studies (https://www.ebi.ac.uk/gwas/variants/rs13007705).

The limitations of our study include the selective use of male mice, preplanned because of known relative resistance of female mice to adenine-induced crystal formation and subsequent CKD (26). There is no reason to expect that ERFE would act differently in female mice, as our previous studies in healthy mice indicate that transgenic augmentation of ERFE has similar effects in both sexes (8).

The transgenic augmentation of ERFE *in utero* impacts embryonic development in TG mice, leading to smaller baseline kidney weight. This developmental effect may lead to underestimation of potential renal benefits of ERFE treatment for CKD-associated anemia. For proof of concept, we chose the transgenic approach because administration of currently available recombinant ERFE may induce an inflammatory response, due to protein aggregation characteristic of the C1q/tumor necrosis factor-related protein family (7). Further technical development will be needed to overcome this limitation or find alternative means of augmenting ERFE. Nonetheless, our findings provide valuable insights into the therapeutic potential of ERFE.

Other approaches specifically targeting the hepcidin-ferroportin pathway in CKD are in various stages of clinical development, and include monoclonal antibodies targeting the positive regulators of hepcidin, including BMP6 (27) and hemojuvelin (28), and monoclonal antibodies to the hepcidin receptor and iron transporter ferroportin (27). In the current study, we employed a common preclinical mouse model of CKD to verify the concept that the endogenous erythroid hormone ERFE can be used therapeutically to ameliorate anemia.

## Materials and Methods

### Mice

The study employs male mice, as CKD in rodents presents a more robust and consistent phenotype in males than in females (29, 30). Mice were maintained in a ventilated rodent-housing system with temperature-controlled environments (22–25°C) with a 12-hour light/dark cycle, and allowed *ad libitum* access to food and water. Unless specified, mice received a standard diet (PicoLab Rodent Diet 20, 5053 Irradiated, 185 ppm iron). Erythroid ERFE-overexpressing transgenic mice were generated as previously detailed(8), and the line with highest ERFE expression (line-H) used in the current study. These mice were maintained on a C57BL/6J background.

To evaluate the therapeutic impact of ERFE on CKD-related complications, wild-type littermates (WT) and erythroid ERFE-overexpressing transgenic (TG) mice were administered a specialized adenine-rich diet with adequate iron content (0.2% adenine, 100 ppm iron; TD.210096, Envigo) for 8 weeks to induce CKD (Figure 1A). To account for the potential confounding variable of incipient chronic iron overload already apparent ∼ 6 weeks of age in TG mice, they were initially placed on an iron-deficient diet (4 ppm iron; TD.80396, Envigo) at weaning for 4 weeks (Supplementary Figure 1G). One cohort of TG mice was analyzed at this point to establish iron and hematological parameters at the onset of the study, and another cohort was switched to the 0.2% adenine diet with adequate iron (100 ppm; TD.210096, Envigo) for an additional 8 weeks (Figure 1F). Age-matched WT mice followed a similar protocol, receiving an iron-adequate diet (100 ppm iron; TD.200065, Envigo) for 4 weeks post-weaning, with one cohort euthanized at this point and another transitioning to the 0.2% adenine-rich, iron-adequate diet for 8 weeks (Figure 1F). Upon completion of these experimental timelines, mice were euthanized under 2.5% isoflurane anesthesia, and sample preparation was conducted as detailed in methods.

### Hematologic parameter and iron-related measurements

Complete blood counts were analyzed using a HemaVet blood analyzer (Drew Scientific). Serum iron and tissue non-heme iron levels were measured through colorimetric quantification following the manufacturer’s instructions (157-30, Sekisui Diagnostics). To reduce variability from regional iron distribution, whole liver, spleen, and kidney tissues were ground in liquid nitrogen before sampling.

### Serum chemistry

Serum hepcidin concentrations were determined by ELISA as previously described (7). Serum urea nitrogen (DIUR-100, BioAssay Systems), and serum creatinine (80350, Cayman Chemical) concentrations were measured by colorimetric quantification. Intact bioactive FGF23 (iFGF23) levels were assessed by ELISA (60-6800, QuidelOrtho). Serum phosphate levels were assessed by colorimetric quantification (830125, ThermoFisher Scientific). Assays were performed following manufacturers’ protocols.

### Cytokine and chemokine measurements

Serum cytokine and chemokine levels were analyzed by the UCLA Integrated Molecular Technologies Core using a multiplex bead-based immunoassay (Millipore Milliplex Cytokine/Chemokine 32-plex Kit on Luminex FlexMap3D), following established protocols (31).

### Histopathology

Kidney tissues were fixed in 10% neutral buffered formalin for 24 hours at room temperature, rinsed twice in distilled water, and stored in 70% ethanol until processing. Tissues were paraffin-embedded, sectioned at 4 μm thickness, and subjected to staining with hematoxylin and eosin (H&E) for morphological analysis or Masson’s trichrome for fibrosis evaluation, performed by the UCLA Translational Pathology Core Laboratory (TPCL). Slides were viewed and imaged using a Nikon Eclipse E600 microscope with SPOT Basic^TM^ (SPOT Imaging) image software.

### Quantitative PCR analysis

Frozen mouse tissues were homogenized in TRIzol Reagent (15596018, ThermoFisher Scientific) and total RNA was extracted using chloroform. Employing a two-step reaction method, 500 ng of total RNA was reverse transcribed into cDNA using the iScript cDNA Synthesis Kit (1708896, Bio-Rad). Quantitative real-time PCR was conducted with 20 ng of cDNA, SsoAdvanced Universal SYBR Green Supermix (172-5272, Bio-Rad) and sequence specific primers (Table 2). Samples were assayed in duplicate with a CFX Connect Real-Time PCR detection system (Bio-Rad). Relative gene expression was normalized to *Hprt* levels. Data are presented as -ΔCt.

### Statistical analysis

Data organization, scientific graphing and statistical significance of differences between experimental groups were performed by using GraphPad Prism (version 10.2.3). Data are presented as individual values representing the mean ± SEM. Statistical differences between groups was determined by unpaired-*t*-test with Welch’s correction (two-tailed). Statistical test, number of animals per group, and *P*-value are indicated in each figure panel and legend. A *P*-value of <0.05 was considered significant.

### Study Approval

All animal studies were approved by the University of California, Los Angeles (UCLA) Institutional Animal Care and Use Committee, and were performed in accordance with the Guide for Care and Use of Laboratory Animals (National Institutes of Health, Bethesda, MD).

## Acknowledgments

This study was supported by the National Institutes of Health through R01 DK126680 (TG, EN) with supplement (JDO), training grant T32 HL072752 (BC), and Ford Foundation Fellowship and Gilliam/Howard Hughes Medical Institute Fellowship (CDC).

