## Supplemental Materials for "Transgenic augmentation of erythroferrone in mice ameliorates anemia in adenine-induced chronic kidney disease"

### Supplemental Figure and Data

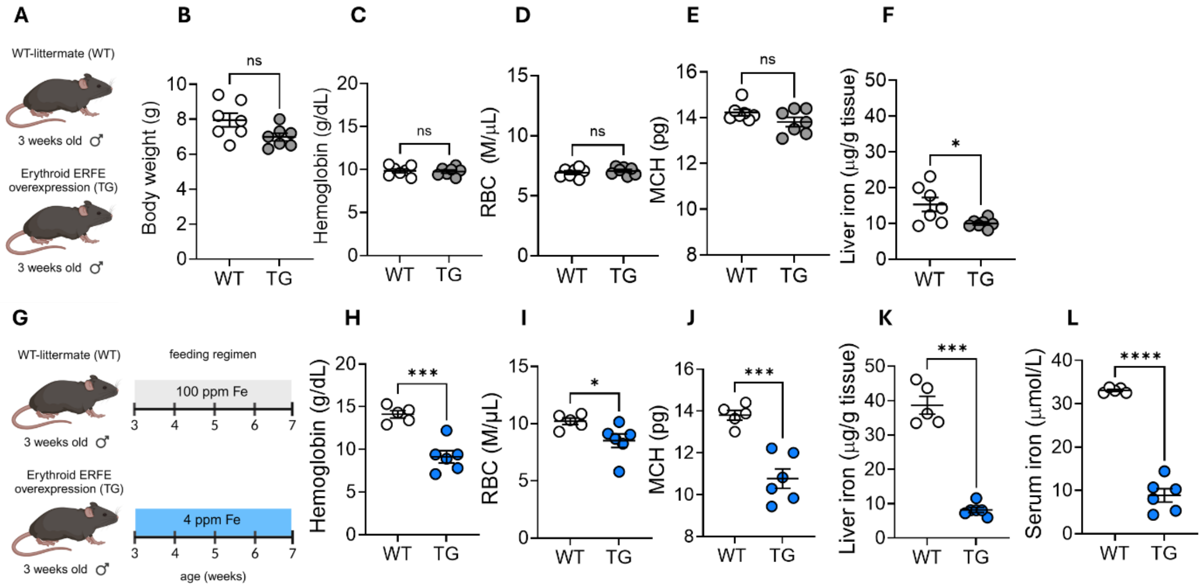


**Figure S1. Experimental design development.** (**A**) WT and TG (n= 7 mice/group) mice at weaning (3 weeks of age), (**B**) body weight, (**C**) hemoglobin concentrations, (**D**) RBC count, (**E**) MCH, (**F**) liver iron concentration (LIC). (**G**) At weaning (3 weeks of age), WT and TG (n=5-6/group) mice were placed on 100 ppm Fe diet and 4 ppm Fe diet respectively, and analyzed after 4 weeks for (**H**) hemoglobin concentration, (**I**) RBC count, (**J**) MCH, (**K**) liver iron concentration (LIC) and (**L**) serum iron concentration. Data are mean ± SEM, analyzed by unpaired-*t*-test with Welch's correction (two-tailed). ***P < .001; *P < .05; ns = non-significant.

**Supplemental Table 1: Body and organ weights of wildtype (WT) and ERFE-TG (TG) mice receiving a 0.2% adenine diet as described in Figure 1F.**

Values are expressed as mean ± SEM. Comparison between groups was performed in form of an

| | WT + 0.2% adenine diet | TG + 0.2% adenine diet |
|---|---|---|
| Starting body weight (g) | 23.7 ± 0.7 | 18.9* ± 0.8 |
| Final body weight (g) | 16.4 ± 0.4 | 16.5 ± 0.4 |
| Wet liver weight (mg) | 691.4 ± 32.4 | 711.5 ± 31.3 |
| Wet spleen weight (mg) | 44.1 ± 3.5 | 54.8* ± 2.4 |
| Wet kidney weight (mg) | 262.5 ± 12.1 | 207.3* ± 13.3 |
| Liver to body weight (%) | 3.78 ± 0.2 | 4.32* ± 0.1 |
| Spleen to body weight (%) | 0.2 ± 0.02 | 0.32* ± 0.02 |
| Kidney to body weight (%) | 1.6 ± 0.1 | 1.2* ± 0.1 |

unpaired-t-test with Welch's correction, N=6/group; *p ≤ 0.05.

**Supplemental Table 2. Oligonucleotides used as sequence specific primers in qPCR analyses.**

| Gene | Species | Orientation | Primer Sequence (5' – 3') |
|---|---|---|---|
| *Acta2* | Mus musculus | Forward<br>Reverse | GTC CCA GAC ATC AGG GAG TAA<br>TCG GAT ACT ACT TCA GCG TCA GGA |
| *Col1a1* | Mus musculus | Forward<br>Reverse | ATG GAT TCC CGT TCG AGT ACG<br>TCA GCT GGA TAG CGA CAT CG |
| *Col3a1* | Mus musculus | Forward<br>Reverse | GAC CAA AAG GTG ATG CTG GAC AG<br>CAA GAC CTC GTG CTC CAG TTA G |
| *Krt20* | Mus musculus | Forward<br>Reverse | CAA CGG ATC GGA CCT GTT TG<br>AGC GCA CTT TTT CTA GGT AGT TT |
| *Fn1* | Mus musculus | Forward<br>Reverse | GAT GTC CGA ACA GCT ATT TAC CA<br>CCT TGC GAC TTC AGC CAC T |
| *Hamp* | Mus musculus | Forward<br>Reverse | GAG CAG CAC CAC CTA TCT CC<br>TTG GTA TCG CAA TGT CTG CC |
| *Hprt* | Mus musculus | Forward<br>Reverse | CTG GTT AAG CAG TAC AGC CCC AA<br>CGA GAG GTC CTT TTC ACC AGC |
| *Il1b* | Mus musculus | Forward<br>Reverse | TGC CAC CTT TTG ACA GTG ATG<br>TGA TGT GCT GCT GCG AGA TT |
| *Il6* | Mus musculus | Forward<br>Reverse | CTC TGG GAA ATC GTG GAA AT<br>CCA GTT TGG TAG CAT CCA TC |
| *Havcr1/Kim1* | Mus musculus | Forward<br>Reverse | CGA GTG GAG ATT CCT GGA TGG<br>GGA CGT GTG GGA ATC TCT GG |
| *Slc5a2/Sglt2* | Mus musculus | Forward<br>Reverse | TGG GCT GGA TAT TTG TCC CGA<br>CAA ACC GCT TCC GCA GAT ACT |
| *Vegfa* | Mus musculus | Forward<br>Reverse | ACA TTG GCT CAC TTC CAG AAA CAC<br>GGT TGG AAC CGG CAT CTT TAT C |
| *Lcn2/Ngal* | Mus musculus | Forward<br>Reverse | GTC CCC ACC GAC CAA TGC<br>ATT GGG TCT CTG CGC ATC C |
| *Gapdh* | Mus musculus | Forward<br>Reverse | CCA ATG TGT CCG TCG TGG ATC T<br>GTT GAA GTC GCA GGA GAC AAC C |
| *Angptl1* | Mus musculus | Forward<br>Reverse | GAC AAT TCA CTC GAA CTT TCC CA<br>CCT TGT TGC CAT CTT CAG CAT T |
| *Saa1* | Mus musculus | Forward<br>Reverse | ACA CCA GCA GGA TGA AGC TAC T<br>GAG CAT GGA AGT ATT TGT CTG AGT |
| *Epo* | Mus musculus | Forward<br>Reverse | TCT ACG TAG CCT CAC TTC ACT<br>ACC CGG AAG AGC TTG CAG AAA |
| *Tgfb* | Mus musculus | Forward<br>Reverse | ATA CGT CAG ACA TTC GGG AAG CAG TG<br>AAT AGT TGG TAT CCA GGG CTC TCC G |
| *Klotho/Kl* | Mus musculus | Forward<br>Reverse | TGA TGT CGT CCA ACA CGT AGG CTT<br>GCA AAG TGC TCA ACT GGC TAA GGT |
| *Tnfa* | Mus musculus | Forward<br>Reverse | CCC TCA CAC TCA GAT CAT CTT CT<br>GCT ACG ACG TGG GCT ACAG |

| | | | |
|---|---|---|---|
| *Id1* | Mus musculus | Forward<br>Reverse | CCT AGC TGT TCG CTG AAG GC<br>CTC CGA CAG ACC AAG TAC CAC |
| *Id2* | Mus musculus | Forward<br>Reverse | ATG AAA GCC TTC AGT CCG GTG<br>AGC AGA CTC ATC GGG TCG T |
| *Id4* | Mus musculus | Forward<br>Reverse | CCG CCC AAC AAG AAA GTC AG<br>CCA GGA TGT AGT CGA TAA CGT G |
| *Smad7* | Mus musculus | Forward<br>Reverse | GGG CTT TCA GAT TCC CAA CTT<br>AGG GCT CTT GGA CAC AGT AGA |
| *Bmp2* | Mus musculus | Forward<br>Reverse | TCA AGC CAA ACA CAA ACA GC<br>AGC CAC AAT CCA GTC ATT CC |
| *Bmp7* | Mus musculus | Forward<br>Reverse | CAT CGT CCA GAC ACT GGT TCA C<br>TCG AAG TAG AGG ACA GAG ATG GC |
| *Slc34a1/NaPi2a* | Mus musculus | Forward<br>Reverse | GGC TCC AAC ATT GGC ACT ACC A<br>ACC ACA GTA GGA TGC CCG AGA T |
| *Slc34a3/NaPi2c* | Mus musculus | Forward<br>Reverse | GCG GTA TTA CCA GCA ACA CCA C<br>TGT CCT CCT CTG GAG ATG CTG A |
| *Cyp27b1* | Mus musculus | Forward<br>Reverse | TTC GGC TTT GGC AAA CGG AGC T<br>GGC TTG ATA GGA AGA GCA CCT G |
| *Cyp24a1* | Mus musculus | Forward<br>Reverse | GCT CCT TCA AAA GGA CAC AGA GG<br>CGC TTG CCA CAC TTT GGT GTT G |
